# *KRAS* signalling in malignant pleural mesothelioma

**DOI:** 10.1101/2020.10.22.350850

**Authors:** Antonia Marazioti, Christophe Blanquart, Anthi C. Krontira, Mario A. A. Pepe, Caroline M. Hackl, Marianthi Iliopoulou, Anne-Sophie Lamort, Ina Koch, Michael Lindner, Rudolph A. Hatz, Darcy E. Wagner, Helen Papadaki, Sophia G. Antimisiaris, Ioannis Psallidas, Magda Spella, Ioanna Giopanou, Ioannis Lilis, Marc Grégoire, Georgios T. Stathopoulos

## Abstract

Malignant pleural mesothelioma (MPM) arises from mesothelial cells lining the pleural cavity of asbestos-exposed individuals and rapidly leads to the development of pleural effusion and death. MPM harbours loss-of-function mutations in genes like *BAP1, NF2, CDKN2A*, and *TP53*, but isolated deletion of these genes alone in mice does not cause MPM and mouse models of the disease are sparse. Here we show that a significant proportion of human MPM harbour point mutations and copy number alterations in the *KRAS* proto-oncogene. These mutations are likely pathogenic, since ectopic expression of mutant *KRAS^G12D^* in the pleural mesothelium of conditional mice causes MPM. Murine MPM cell lines derived from these tumours carry the initiating *KRAS^G12D^* lesions, secondary *Bap1* alterations, and human MPM-like gene expression profiles. Moreover, they are transplantable and actionable by KRAS inhibition. Our results indicate that *KRAS* mutations likely play an important and underestimated role in MPM, which warrants further exploration.

---

Malignant mesothelioma annually kills up to forty persons per million population worldwide^1,2^. It most commonly arises from the mesothelium of the pleural cavities that lines the lungs (visceral pleura) and the interior chest wall (parietal pleura) and only occasionally from the peritoneal mesothelium. Human MPM is mainly caused by inhaled asbestos exposure, but nanofibers can also cause mesothelioma in rodents and possibly in humans^2–4^. MPM commonly manifests with malignant pleural effusions (MPE), i.e., exudative fluid accumulation that causes chest pain and dyspnea^2^, and is histologically classified into epithelioid, sarcomatoid, or biphasic subtypes^5,6^. The disease progresses relentlessly despite contemporary combination therapies, with a median survival of mere 9-18 months^7,8^. The clinicopathologic manifestation of MPM at diagnosis impacts patient survival, with advanced stage, sarcomatoid histologic subtype, poor physical performance status, and the presence of MPE being dismal prognostic indices^9–12^.

Multiple comprehensive analyses of MPM genomes identified a mosaic mutational landscape characterized by widespread loss-of-function of tumour suppressor genes (*BAP1, NF2, CDKN2A, TP53, TSC1*, etc), sporadic gain-of-function of protooncogenes (*PIK3CA, EGFR, KRAS, NRAS, HRAS, BRAF*, etc), and inconclusive addiction/exclusion patterns thereof^13–23^. Interestingly, oncogene mutations along the RAS pathway were detected more frequently by targeted sequencing approaches compared with massive parallel sequencing^13–23^. In addition, *NF2* mutations that cause persistent RAS signalling^24^, as well as *BAP1* and *CDKN2A* mutations that are functionally related with *TP53* loss-of-function^25–27^, are very common in MPM^13–23^. In addition to clinicopathologic presentation, MPM mutations also impact prognosis, with *TP53* loss-of-function occurring exclusively in sarcomatoid MPM and portending poor survival^14,19^.

There is an unmet clinical need for mouse models that recapitulate the mutation spectrum and clinicopathologic manifestations of human MPM. In this regard, MPM cell lines for transplantable models, asbestos-induced mouse models, as well as genetic models of the disease are characterised by scarcity, limited availability, and significant difficulty of implementation^28–34^. Interestingly, standalone mesothelial loss-of-function of *BAP1, NF2, CDKN2A, TP53*, and *TSC1* is not sufficient to cause MPM in mice^31–34^, rendering the true drivers of the disease resistant to functional validation. Moreover, faithful models of MPM are urgently needed, as most existing studies have focused on the rare peritoneal disease^32–34^ and only one elegant study targeted *NF2/CDKN2A/TP53* deletions to the pleural mesothelium^31^. Even the latter model, however, does not yield MPE, which are observed in 90% of human MPM^2^.

Based on our previous observation of a *Kras^G12C^* mutation in an asbestos-induced murine MPM cell line^35^ and in published work that showed RAS pathway activation in MPM^36^, we hypothesized that *KRAS* mutations are involved in MPM development. Indeed, here we employ sensitive methods to discover point mutations and amplifications in a significant proportion of MPM tumours from our clinical cohorts. We further show that targeting oncogenic *KRAS*^G12D^ alone to the murine pleural mesothelium causes MPM and, when combined with *Trp53* deletion, triggers aggressive MPM with MPE. Murine MPM are shown to carry the initiating *KRAS*^G12D^ mutations, to harbour *Bap1* inactivating mutations, to be transmissible to naïve mice, and to resemble the molecular signatures of human MPM. Hence, *KRAS* mutations are implicated in MPM pathobiology, the contributions of *TP53* in shaping the disease’s manifestations is elucidated, and new mouse models are provided for the study of MPM biology and therapy.

## RESULTS

### Identification of *KRAS* mutations in human MPM patients

We first employed digital droplet polymerase chain reaction (ddPCR) in order to detect *KRAS* mutations in MPE cell pellets of 12 patients with MPM and 14 with lung adenocarcinoma (LUAD) from Munich, Germany^37,38^. The assay was designed for detection of down to 1:20000 mutant copies using EKVX (*KRAS*-wild type, *KRAS*^WT^) and A549 (*KRAS*^G12S^) cells as negative and positive controls, respectively. We found *KRAS* mutations in three patients from each group, with two patients per group displaying low mutant copy numbers (< 1:200) that would be likely missed by other techniques (Figs. 1a,b). Two patients with MPM and one with LUAD had codon 12/13 mutations, while one patient with MPM and two with LUAD had codon 61 mutations. We next used Affymetrix CytoScanHD Arrays to identify copy number alterations of the *KRAS* locus in a cohort of primary MPM cell lines from Nantes, France (GEO dataset GSE134349)^39,40^. We found gains in ten and losses in two of 32 MPM cell lines examined, which was statistically significantly more that what was reported in two previous studies of MPM tumours^22,41^ (Fig. 1c). Finally, we calculated combined RAS (*KRAS, EGFR, PIK3CA*) and TP53 (*TP53, STK11*) pathway mutation rates in a selected set of human MPM genomic studies^13–21^, and found each pathway to account for 10% of the total mutations observed (Fig. 1d). These results show that *KRAS* point mutations and gains are present in a subset of MPM.

**Figure 1.**
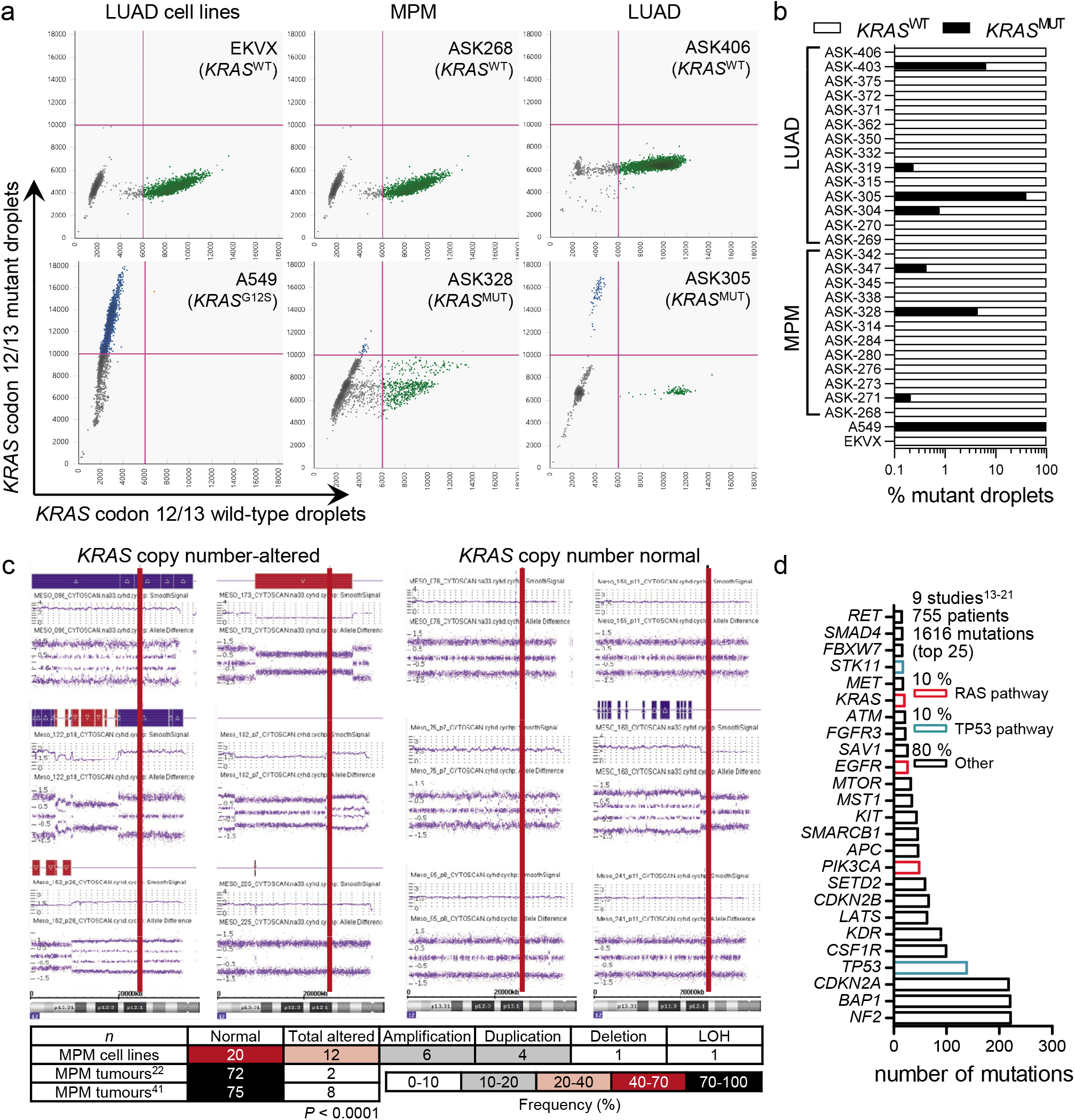
KRAS mutations in human malignant pleural mesothelioma (MPM). **a,b,** Digital droplet polymerase chain reaction (ddPCR) for codon 12/13 and codon 61 *KRAS* mutations in MPE cell pellets of 12 patients with MPM and 14 with lung adenocarcinoma (LUAD) from Munich, Germany^37,38^, as well as EKVX (*KRAS*-wild type, *KRAS*^WT^) and A549 (*KRAS*^G12S^) cells. Shown are representative dotplots (a) and data summary of mutant KRAS copies percentage (b). **c,** Representative results (images) and results summary (table) from Affymetrix CytoScanHD Arrays of 32 primary MPM cell lines from Nantes, France (GEO dataset GSE134349)^39,40^, compared with published results from two previous studies of MPM tumours^22,41^. Red lines denote the KRAS locus on chromosome 12p12.1. Colour code denotes frequency. *P*, probability, χ^2^ test. **d,** Mutation rate of the top twenty-five mutated genes of human MPM generated by combining data from ten published studies^13–21^. Red and blue columns indicate RAS and TP53 pathway genes. RAS, rat sarcoma; TP53, tumour-related protein 53.

### MPM in mice expressing mesothelial-targeted *KRAS*^G12D^

To functionally validate *KRAS* mutations in MPM, we targeted transgenes to mesothelial surfaces using type 5 adenoviral vectors (Ad). For this, *mT/mG* CRE-reporter mice that switch from somatic cell membranous tomato (mT) to green fluorescent protein (mG) expression upon CRE-mediated recombination^42^ received 5 x 10^8^ plaque-forming units (PFU) intrapleural Ad encoding *Melanotus* luciferase (Ad-*Luc*) or Ad-*Cre* followed by serial bioluminescence imaging. Ad-*Luc*-treated mice developed intense bilateral chest light emission (mice lack mediastinal separations)^43^ that peaked at 4-7 and subsided by 14 days post-injection (Fig. 2a). At this time-point, when transient Ad-*Luc* expression ceased and therefore maximal Ad-*Cre*-mediated recombination was achieved, Ad-*Cre*-treated mice displayed widespread recombination of the pleural mesothelium even in contralateral pleural fissures, but not of the lungs, chest wall, or pleural immune cells (Figs. 2b–e). Similar results were obtained from intraperitoneal 5 x 10^8^ PFU *Ad-Cre*-treated *mT/mG* mice after two weeks (Fig. 2f). Importantly, Ad-*Cre* did not cause inflammation, as evident by imaging and cellular analyses of luminescent bone marrow chimeras used as real-time myeloid tracers (Figs. 3a,b)^35^. These results show that intraserosal Ad-*Cre* treatment efficiently and specifically recombines mesothelial surfaces *in vivo*. To test whether oncogenic *KRAS* can cause MPM, wild-type (*Wt*) mice and mice carrying conditional *KRAS^G12D^* and/or *Trp53f/f* alleles expressed or deleted, respectively, upon CRE-mediated recombination^44–46^ received 5 x 10^8^ PFU intrapleural Ad-*Cre*. *Wt, Trp53f/Wt*, and *Trp53f/f* mice survived up to 16 months post-Ad without clinical or pathologic disease manifestations (median survival undefined). In contrast, *KRAS^G12D^* mice developed cachexia and succumbed by 6-12 months post-injection [median(95% CI) survival = 339(285-379) days; *P* = 0.005 compared with controls, log-rank test; Fig. 4a]. At necropsy, no pleural fluid or inflammatory cell accumulation was evident, but diffuse visceral and parietal pleural nodular and peel-like lesions were found in all mice. These lesions expressed proliferating cell nuclear antigen (PCNA) unlike the normal pleura and were diagnosed by a board-certified pathologist as epithelioid MPM (Figs. 4b–d). In addition, chimeric *KRAS^G12D^* recipients adoptively transplanted with luminescent bone marrow revealed an early pleural inflammatory infiltrate composed of CD11b+Gr1+ myeloid cells at 7-14 days post-Ad-*Cre* (Figs. 2a,b), similar to what is observed after pleural asbestos instillation^3^. The phenotype of intrapleural Ad-*Cre*-injected *KRAS^G12D^;Trp53f/f* mice was fulminant, with respiratory and locomotor distress and retracted body posture culminating in death by 3-6 weeks post-Ad-*Cre* [median(95% CI) survival = 41(38-73) days; *P* < 0.001 compared with any other genotype, log-rank test; Fig. 4a]. Examination of the thorax revealed massive MPE in most and visceral/parietal pleural tumours in all mice, which invaded the lungs, chest wall, and mediastinum and uniformly presented as PCNA+ biphasic MPM with mixed sarcomatoid/epithelioid features (Figs. 4b–f). Effusions were bloody but non-coagulating, contained abundant cancer and inflammatory cells, and had low pH and glucose and high protein, VEGF, and lactate dehydrogenase levels (Fig. 4g), resembling effusions of human advanced MPM^2^ and of *C57BL/6* mice injected with *KRAS^G12C^*-mutant AE17 mesothelioma cells^35^. *KRAS^G12D^;Trp53f/Wt* mice displayed an intermediate phenotype [median(95% CI) survival = 118(97-160) days; *P* < 0.003 compared with any other genotype, log-rank test], biphasic histology, and a single MPE occurrence (Figs. 4a–f). *Wt, Trp53f/f*, and *KRAS^G12D^;Trp53f/f* mice also received 5 x 10^8^ PFU intraperitoneal Ad-*Cre*. Again, *Wt* and *Trp53f/f* mice displayed unlimited survival without signs of disease (median survival undefined), but *KRAS^G12D^;Trp53f/f* mice developed abdominal swelling and succumbed by 2-5 months post-Ad-*Cre* [median(95% CI) survival = 95(60-123) days; *P* < 0.001 compared with controls, log-rank test; Fig. 5a]. At necropsy, nodular and diffuse tumors throughout the abdominal cavity and loculated ascites with features similar to MPM with MPE were detected (Figs. 5b–d). These results show that pleural mesothelial-targeted *KRAS^G12D^* causes epithelioid MPM in mice. Furthermore, that standalone *TP53* loss does not trigger MPM, but cooperates with mutant *KRAS* to accelerate mesothelioma development, to promote biphasic histology, and to precipitate effusion formation.

**Figure 2.**
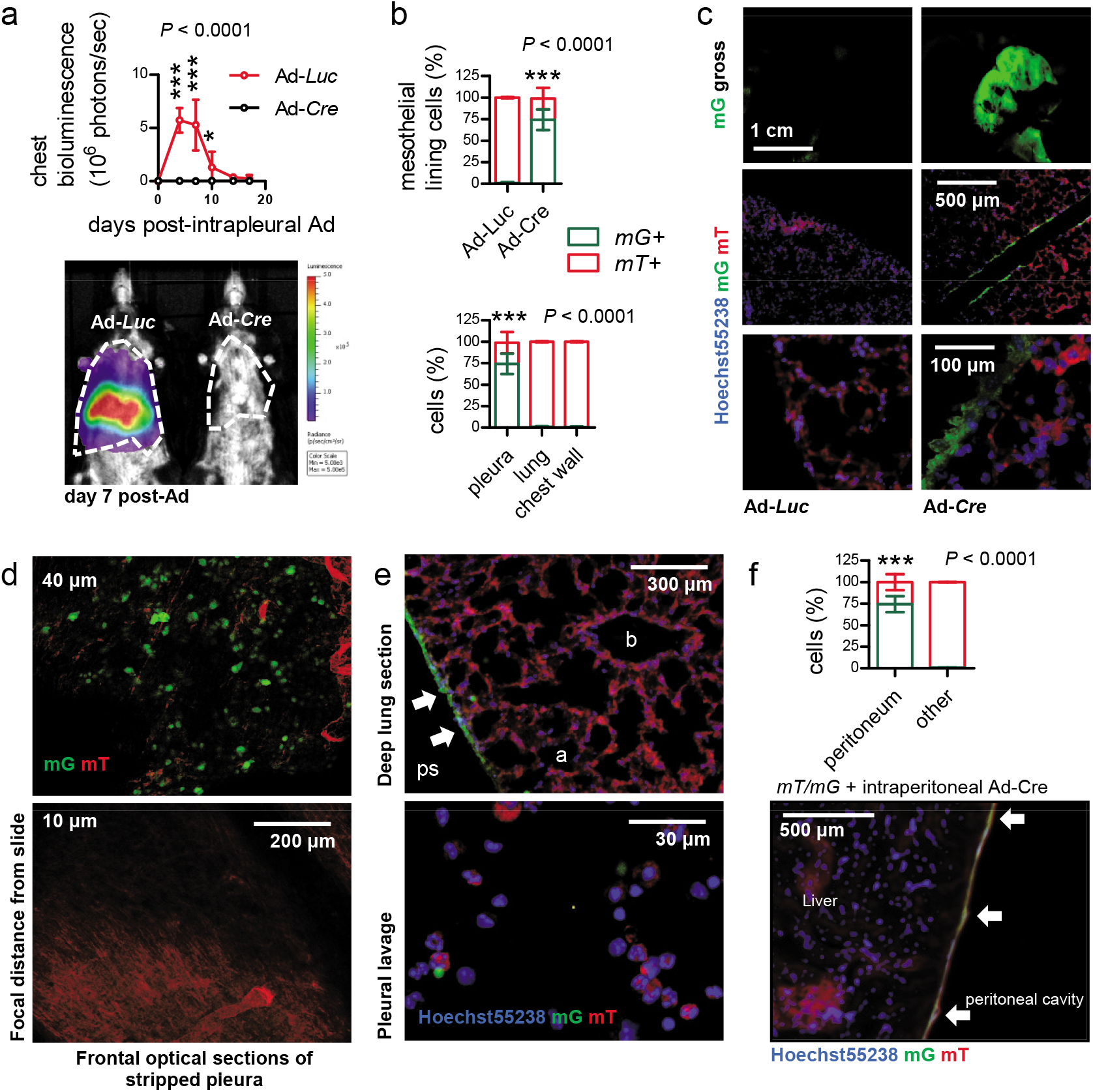
Adenoviral-mediated mesothelial recombination. Dual-fluorescent *mT/mG* CRE-reporter mice (*C57BL/6* background) received 5 x 10^8^ PFU intrapleural (a-e) or intraperitoneal (f) Ad-*Luc* or Ad-*Cre* and were serially imaged for bioluminescence. **a,** Data summary of chest light emission (left; *n* = 5 mice/group) and representative bioluminescence images (right). Note cessation of transient Ad-*Luc* expression by day 14. **b–e,** Data summary of mG+ and mT+ mesothelial, lung, and chest wall cell percentage (*n* = 10 mice/group (b), representative macroscopic (top) and microscopic (bottom) fluorescent images (c), optical frontal sections of stripped parietal pleura (d), and deep lung sections (e, top) and fluorescent image of pleural lavage cells (e, bottom). **(f)** Data summary of mG+ and mT+ mesothelial and deeper located (other) abdominal cell percentage (*n* = 10 mice/group) and representative merged microscopic fluorescent image of peritoneal surface mesothelium showing *Cre*-recombined mesothelium (arrows). Data are presented as mean ± 95% confidence interval. *P*, overall probability, two-way ANOVA. * and ***: *P* < 0.05 and *P* < 0.001 for comparison between groups at the indicated time-points (a) or with all other groups by Bonferroni post-tests. Ad, adenovirus; PFU, plaque-forming units; *Luc*, luciferase gene; *Cre*, CRE recombinase gene; *mT*, membranous tomato red; *mG*, membranous green fluorescent protein; ANOVA, analysis of variance.

**Figure 3.**
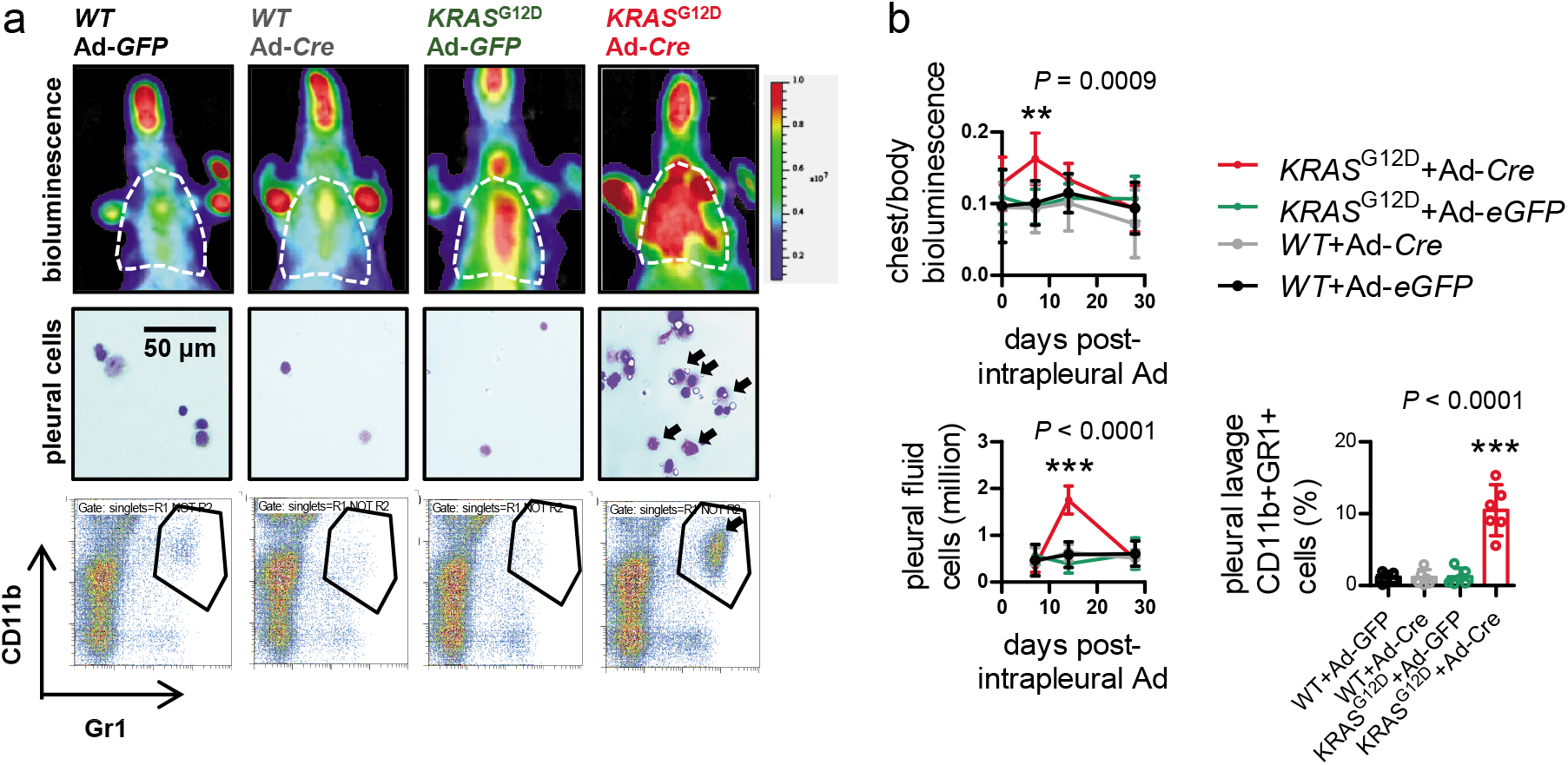
Pleural mesothelial *KRAS^G12D^* expression causes inflammation. Wildtype (*WT*) and *KRAS^GD^* mice were lethally irradiated (1100 Rad) and received same-day bone marrow transfer of ten million bone marrow cells from ubiquitously luminescent *CAG.Luc.eGFP* donors (all on the *C57BL/6* strain)^35,65^. After one month required for bone marrow reconstitution, chimeras received 5 x 10^8^ PFU intrapleural Ad vectors, were longitudinally imaged for bioluminescence, and were sacrificed for pleural lavage cell analysis. **a,** Representative chest bioluminescence images taken two weeks post-Ad (top), pleural lavage cytocentrifugal specimens stained with May-Gruenwald-Giemsa (middle), and dotplots of CD11b and Gr1 expression by flow cytometry (bottom). **b,** Summary of longitudinal chest light emission and total pleural cell number (dotplots), legend to dotplots, as well as of CD11b+Gr1+ pleural cells at day 14 post-Ad (bar graph) shown as mean ± 95% confidence interval (*n* = 5-6 mice/data-point). *P*, overall probability, one-way (bar graph) or two-way (dotplots) ANOVA. ** and ***: *P* < 0.01 and *P* < 0.001, respectively, for Ad-*Cre*-treated *KRAS*^G12D^ mice compared with all other groups by Bonferroni post-tests. *WT*, wildtype; *KRAS^G12D^*, Lox-STOP-Lox.KRAS^G12D^; *CAG.Luc.eGFP*, ubiquitously luminescent mice; Ad, adenovirus type 5; PFU, plaque-forming units; *Cre*, CRE recombinase gene; GFP, green fluorescent protein; ANOVA, analysis of variance.

**Figure 4.**
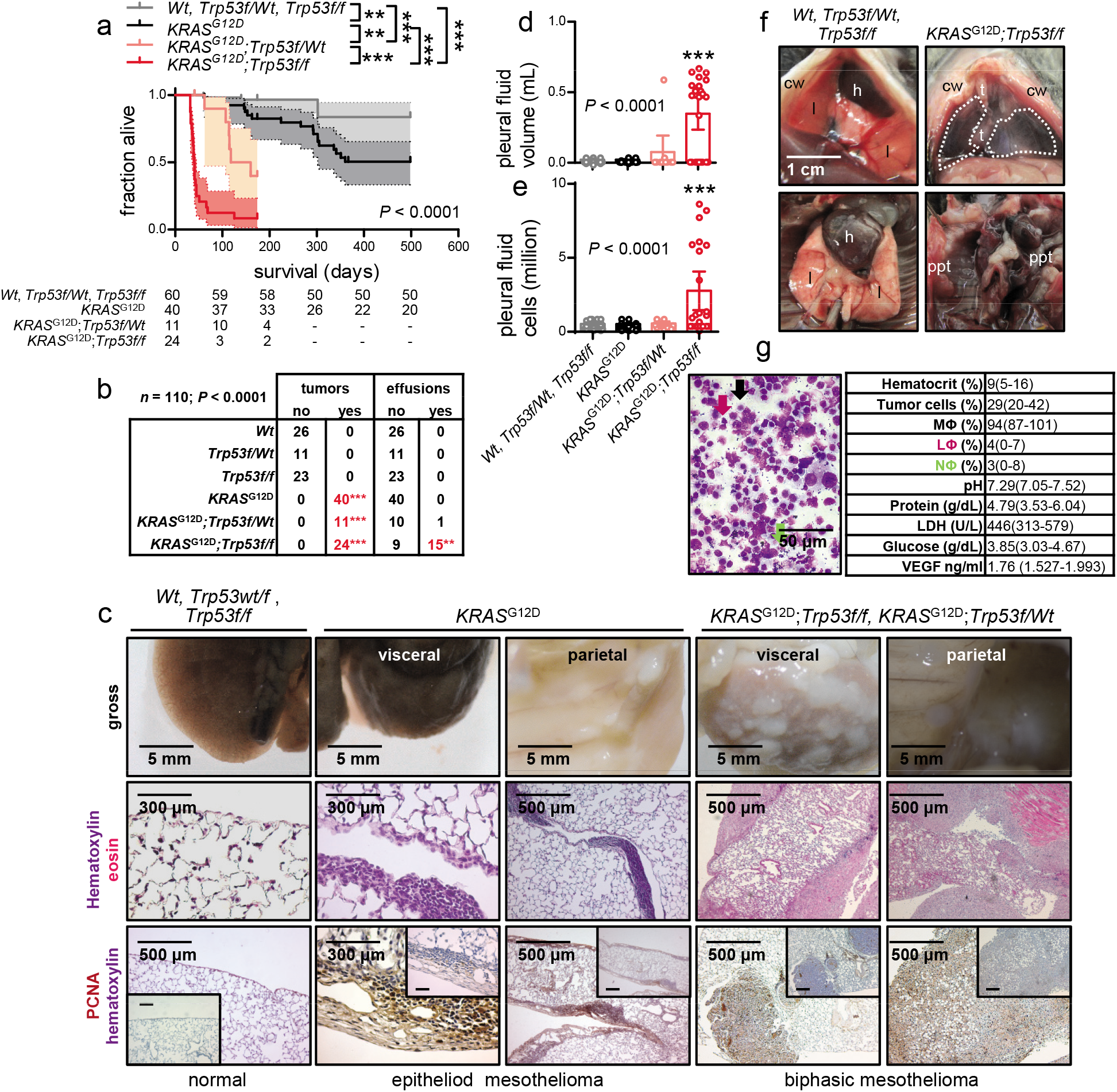
Malignant pleural mesotheliomas and effusions of mice with pleural mesothelial-targeted oncogenic *KRAS^G12D^* and/or *Trp53* deletion. Wild-type (*Wt*), *KRAS^G120^*, and *Trp53f/f* mice (all *C57BL/6*) were intercrossed and all possible offspring genotypes received 5 x 10^8^ PFU intrapleural Ad-*Cre* [n is given in survival table in (a)]. **a,** Kaplan-Meier survival plot. **b,** Incidence of pleural tumours and effusions. **c,** Gross macroscopic and microscopic images of visceral and parietal tumours stained with hematoxylin and eosin or PCNA (n is given in b). **d,e,** Data summary of pleural effusion volume and nucleated cells [n is given in table in (b)]. **f,** Representative photographs of the thorax before (top) and after (bottom) chest opening (t, tumours; l, lungs; cw, chest wall; h, heart; dashed lines, effusion; ppt, parietal pleural tumors). **g,** Representative May-Gruenwald-Giemsa-stained pleural fluid cytocentrifugal specimen from a *KRAS*^G12D^;*Trp53f/f* mouse showing macrophages (MΦ, black arrows), lymphocytes (LΦ, purple arrows), and neutrophils (NΦ, green arrows) and summary of cellular and biochemical features of effusions of *KRAS*^G12D^;*Trp53f/f* mice (*n* = 10). Data are presented as survival fraction ± 95% confidence interval (shaded areas) and number of surviving mice (a), number (*n*) of mice (b), or mean ± 95% confidence interval (d, f). *P*, overall probability, log-rank test (a), χ^2^ test (b), or one-way ANOVA (d). ** and ***: *P* < 0.01 and *P* < 0.001 for comparison indicated (a), with the top-three groups (b), or with all other groups (d) by log-rank test (a), Fischer’s exact test (b), or Bonferroni post-tests (d). *Wt*, wild-type; *KRAS^G12D^*, Lox-STOP-Lox. *KRAS*^G12D^; *Trp53f/f*, conditional *Trp53*-deleted; Ad, adenovirus type 5; PFU, plaque-forming units; *Cre*, CRE recombinase gene; PCNA, proliferating cell nuclear antigen; LDH, lactate dehydrogenase; ANOVA, analysis of variance.

**Figure 5.**
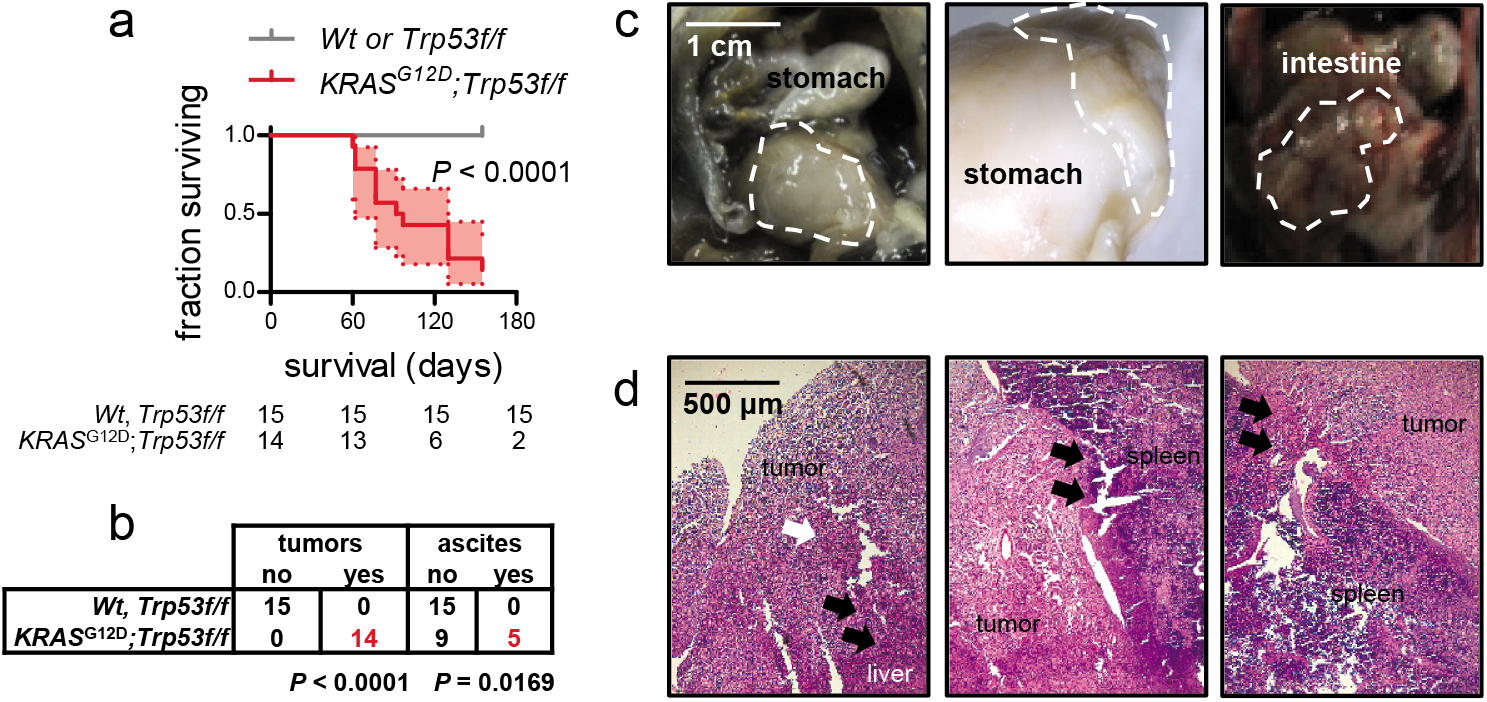
Malignant peritoneal mesothelioma of *KRAS^G12D^;Trp53f/f* mice. Wild-type (*Wt*), *Trp53f/f*, and *KRAS^G12D^;Trp53f/f* mice (all *C57BL/6*) received 5 x 10^8^ PFU intraperitoneal Ad-*Cre* and were harvested when moribund. **a,** Kaplan-Meier survival estimates ± 95% confidence interval (shaded areas) and numbers of surviving mice. **b,** Tumour and ascites incidence table. **c,** Representative macroscopic images of peritoneal tumours (dashed outlines). **d,** Representative hematoxylin and eosin-stained tissue sections of peritoneal tumours. *P*, probability, log-rank test (a) or χ^2^ test (b). *Wt*, wild-type; *KRAS^G12D^*, Lox-STOP-Lox.*KRAS^G12D^*; *Trp53f/f*, conditional *Trp53*-deleted; Ad, adenovirus type 5; PFU, plaque-forming units; *Cre*, CRE recombinase gene.

### Molecular pathology of murine MPM

To corroborate that our mice had mesothelioma and not pleural spread of lung adenocarcinoma^44^, immunostaining for specific markers of both tumour types was performed^5,31,47^ based on expert guidelines for distinguishing MPM from LUAD^6^. In parallel, LUAD of intratracheal Ad-*Cre*-treated (5 x 10^8^ PFU) *KRAS^G12D^* and of urethane-treated mice were examined^48^. MPM expressed calretinin, podoplanin, cytokeratin 5/6, and osteopontin, but not surfactant protein C, in contrast with LUAD that expressed some of these markers and surfactant protein C (Fig. 6), supporting that our tumours are indeed MPM.

**Figure 6.**
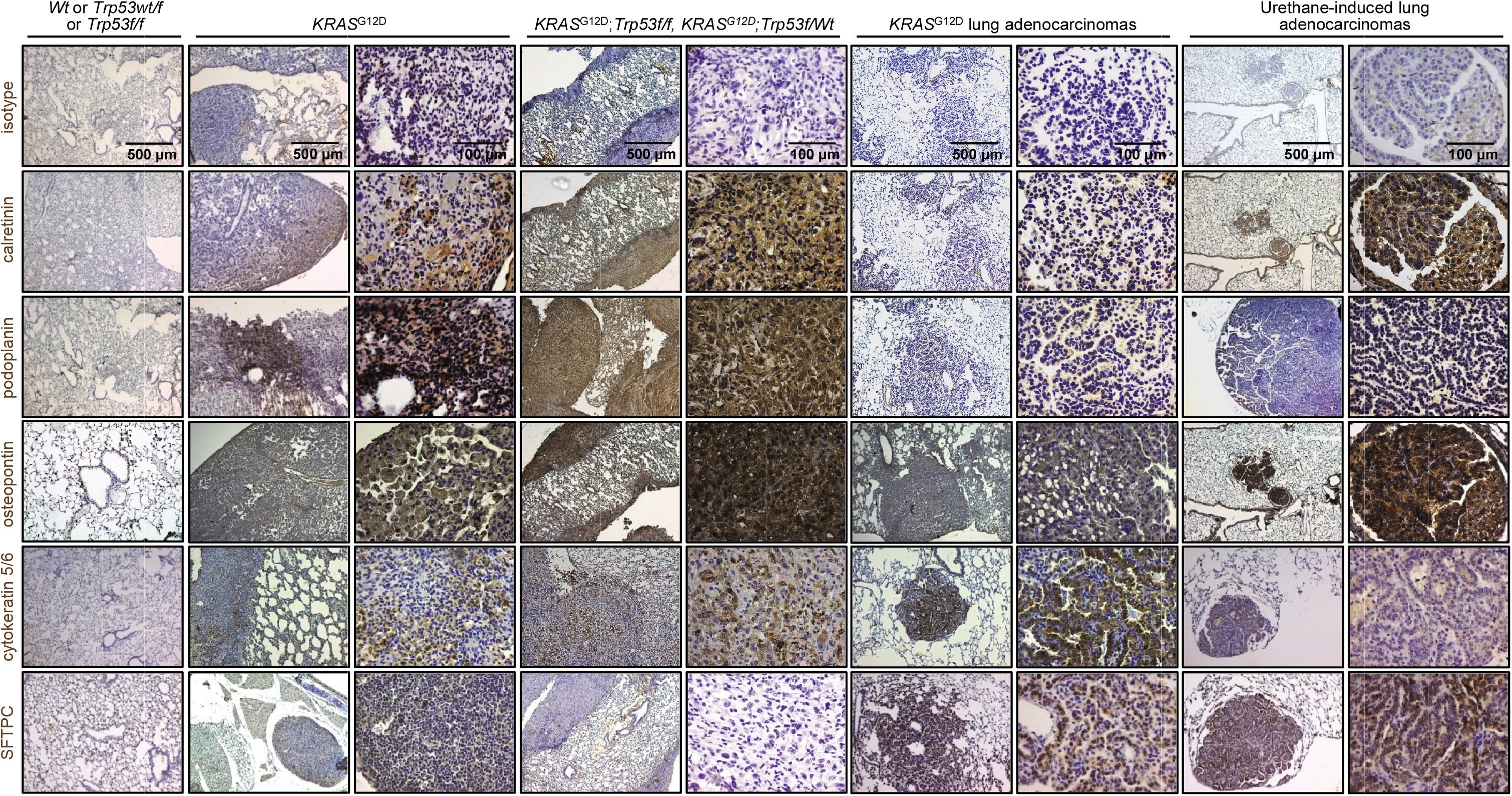
Molecular phenotyping of murine mesothelioma. Wild-type (*Wt*), *KRAS^G12D^*, and *Trp53f/f* mice were intercrossed, and all possible offspring genotypes received 5 x 10^8^ PFU intrapleural or intratracheal Ad-*Cre* and were sacrificed when moribund. In parallel, *C57BL/6* mice received ten consecutive weekly intraperitoneal injections of 1 g/Kg urethane and were sacrificed after six months. Representative images of immunoreactivity of tissue sections of pleural and pulmonary tissues and tumours from these mice for different markers of human mesothelioma and lung adenocarcinoma (*n* = 10 mice/group were analysed for each marker). Brown colour indicates immunoreactivity and blue colour nuclear hematoxylin counterstaining. Note the strong expression of calretinin, podoplanin, osteopontin, and cytokeratin 5/6 and the absence of expression of surfactant protein C (SFTPC) in murine mesotheliomas. Note also the strong expression of osteopontin and SFTPC, and the varying expression of calretinin, podoplanin, and cytokeratin 5/6 in murine lung adenocarcinomas.

### Murine MPM cell lines with *KRAS*^G12D^, Trp53 loss, and *Bap1* mutations and a human-like transcriptome

We subsequently isolated three different MPM cell lines from Ad-*Cre*-treated *KRAS^G12D^;Trp53f/f* mice (KPM1-3). KPM cells displayed anoikis, spindle-shaped morphology, and rapid growth in minimal-supplemented media and in soft agar, were tumorigenic when injected subcutaneously into the flank of *C57BL/6* mice, and carried the original *KRAS^G12D^/Trp53* lesions (Figs. 7a–f). KPM cells and their parental tumours featured enhanced *Cdkn2a* but not *Nf2* expression (Figs. 7e–g, Supplementary Fig. 1), consistent with *TP53*-mediated repression of *BRCA1* and *CDKN2A* expression^25,27^. RNA sequencing of KPM cells (GEO dataset GSE94415) revealed that they carry the pathogenic *KRAS^G12D^/Trp53* lesions, but also multiple single nucleotide variants in exon 6 and insertions in exon 11 of *Bap1*, all of which were validated by Sanger sequencing and immunohistochemistry (Figs. 8a–c). These results support that the three KPM cell lines display different patterns of stochastic *Bap1* mutations, summarized in Fig. 8d.

**Figure 7.**
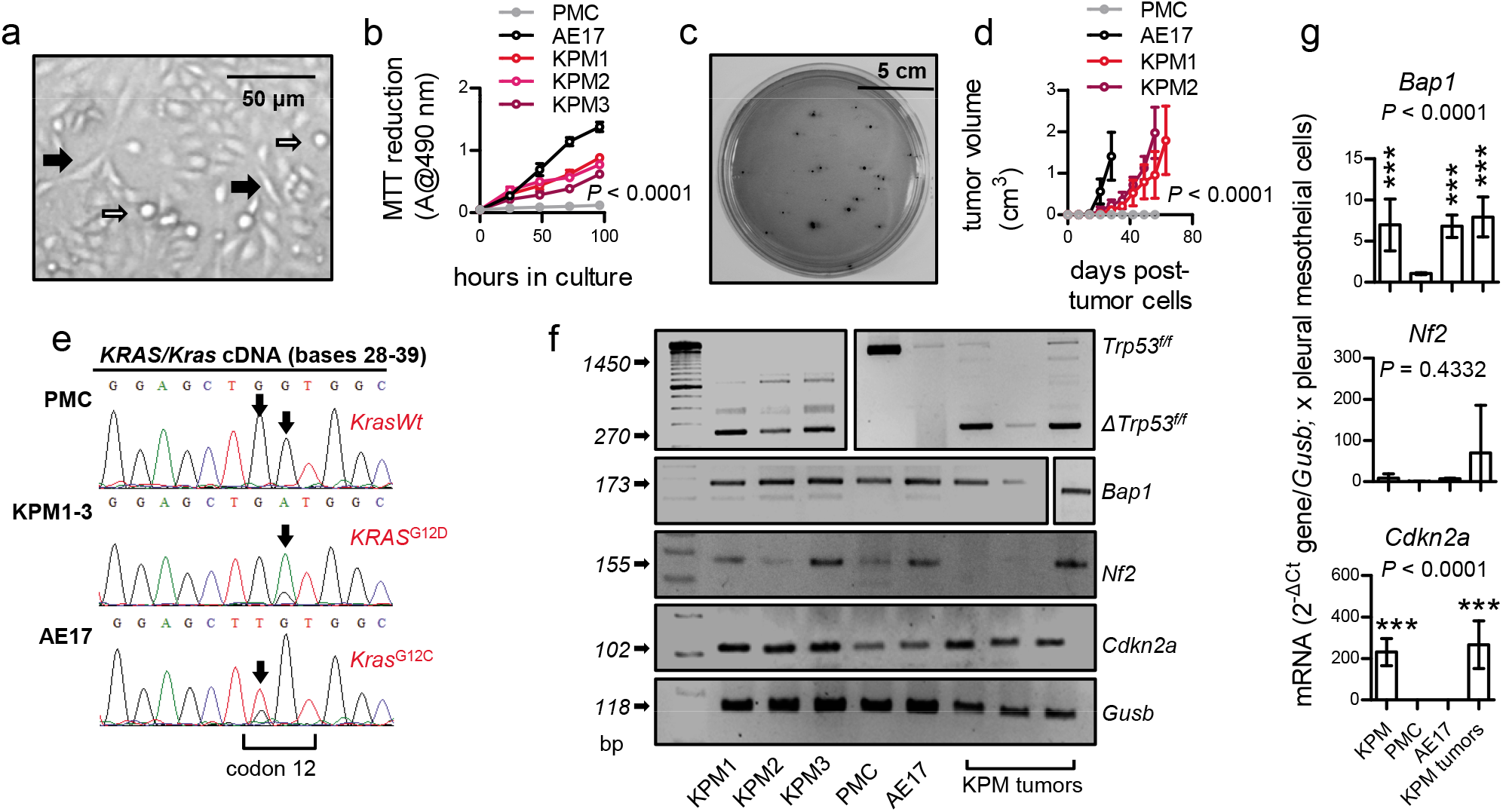
Transplantable *KRAS/TP53*-mutant murine mesothelioma (KPM) cell lines. *KRAS^G12D^;Trp53f/f* pleural mesothelioma (KPM), pleural mesothelial (PMC), and asbestos-induced AE17 mesothelioma cells (all from *C57BL/6* mice) were analyzed. **a,** KPM cell culture showing anoikis (white arrows) and spindle-shaped morphology (black arrows). **b,** *In vitro* MTT reduction (2 x 10^4^ cells/well; *n* = 3 independent experiments). **c,** Representative colonies of KPM1 cells (7.5 x 10^3^ cells/vessel) seeded on a soft agar-containing 60 mm Petri dish and stained with crystal violet after a month (*n* = 3/group). **d,** *In vivo* subcutaneous tumour growth after injection of 10^6^ *cells/C57BL/6* mouse (*n* = 10/group). **e,** *KRAS/Kras* mRNA Sanger sequencing shows *Wt Kras* of PMC, *KRAS*^G12D^ of KPM, and *Kras*^G12C^ of AE17 cells (arrows). **f,g,** RT-PCR (f) and qPCR (g) of KPM cells and parental tumours show *Trp53f/f* allele deletion (*Δ*) and *Bap1* and *Cdkn2a* overexpression compared with PMC. Data are presented as mean ± 95% confidence interval. *P*, overall probability, two-way (c, d) or one-way (g) analysis of variance.

**Figure 8.**
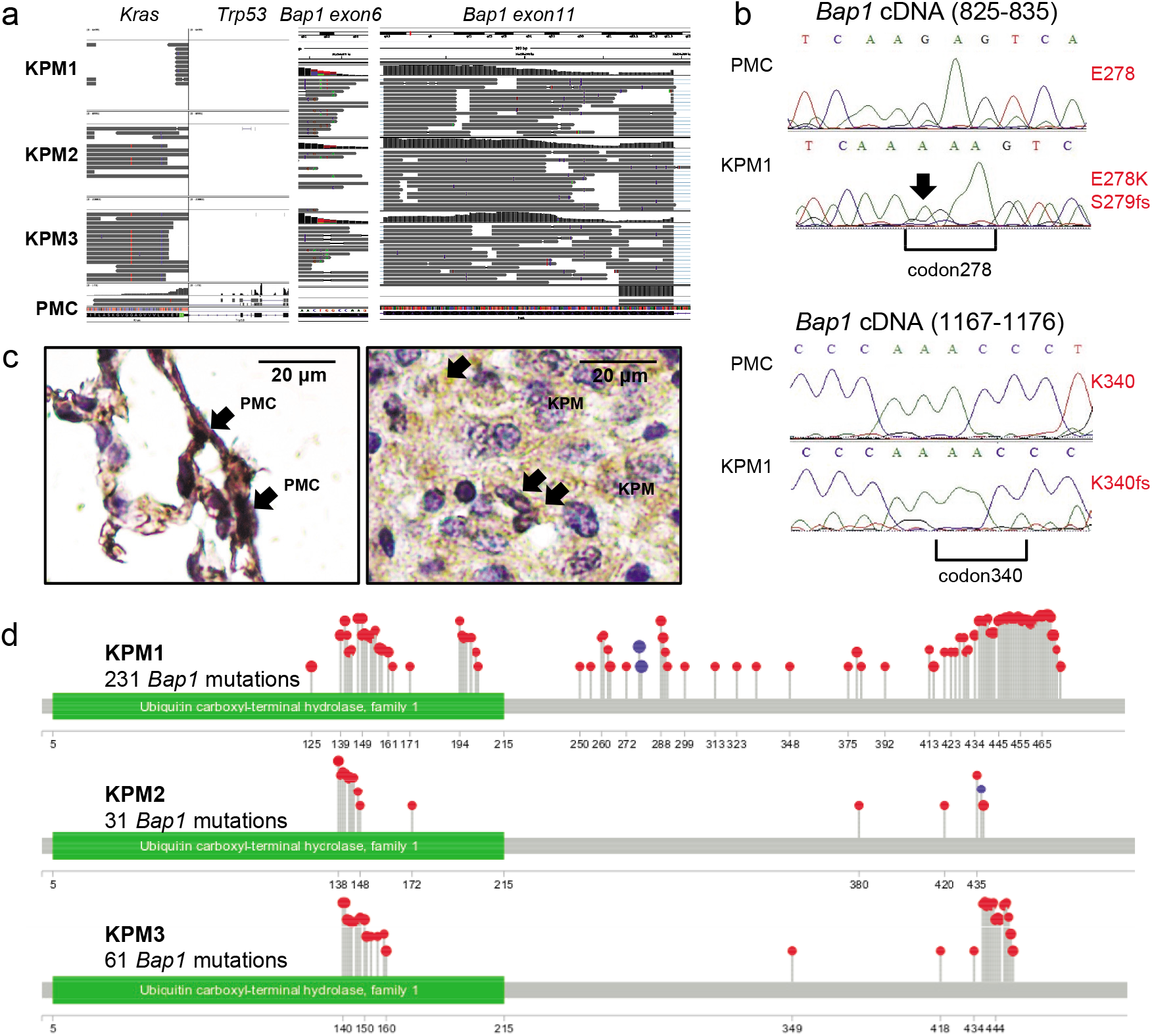
Bap1 mutations of KPM cells. *KRAS^G12D^;Trp53f/f* pleural mesothelioma (KPM) and pleural mesothelial cells (PMC) were analyzed by RNA sequencing (GEO dataset GSE94415), Sanger sequencing for *Bap1*, and immunohistochemistry for BAP1 protein expression. **a,** Coverage and alignment plot from RNA sequencing. Alignments are represented as gray polygons with reads mismatching the reference indicated by colour. Loci with a large percentage of mismatches relative to the reference are flagged in the coverage plot as colour-coded bars. Alignments with inferred small insertions or deletions are represented with vertical or horizontal bars, respectively. **b,** *Bap1* mRNA Sanger sequencing shows a G>A transition at c.829 that generates a missense mutation in codon E278K, a single nucleotide insertion in position c.831 with a consequent frameshift mutation in codon S279insA and a single nucleotide insertion resulting to a frameshift mutation in codon K340insA at c.1072. **c,** Representative immohistochemical images of BAP1 immunoreactivity (brown) of lungs with normal PMC and mouse tumours caused by transplanted KPM cells counterstained with hematoxylin (blue). **d,** Lollipop plot for each KPM cell line visualizing all *Bap1* mutations detected.

### Transplantable and actionable MPM in mice injected with primary KPM cells

Finally, 2 x 10^5^ pleural-delivered KPM cells could inflict to naïve *C57BL/6* mice secondary disease identical to primary MPM of *KRAS^G12D^;Trp53f/f* mice in terms of manifestation, pathology, cytology, and biochemistry (Figs. 9a–f), fulfilling modified Koch’s postulates^49^. To determine the potential efficacy of KRAS inhibition against murine MPM, *C57BL/6* mice received pleural KPM1 cells, followed by a single intrapleural injection of liposomal-encapsulated deltarasin (15 mg.Kg^−1^) or empty liposomes on day nine post-tumour cells, in order to allow initial tumour implantation in the pleural space^35^. At day 19 after pleural injection of KPM1 cells, deltarasin-treated *C57BL/6* mice developed fewer and smaller MPE compared with controls (Fig. 9g). These results collectively show that our murine MPM are indeed malignant, originate from recombined mesothelial cells, and cause transplantable disease that can be used for drug testing. Furthermore, the data support hypotheses on a tumour-initiating role for *KRAS* in MPM (Fig. 9h). RNA sequencing of KPM cells comparative to normal pleural mesothelial cells (PMC) revealed a distinctive transcriptomic signature that included classic mesothelioma markers (*Msln, Spp1, Efemp1, Pdpn, Wt1*) as well as new candidate mesothelioma genes (Figs. 10a–c; Supplementary Table S1). A human 150-gene mesothelioma signature derived from a cohort of 113 patients via comparison of MPM against multiple other malignancies^50^ was highly enriched in our KPM cell line signature (Fig. 10d). These data indicate that murine *KRAS/TP53*-driven MPM present a gene expression profile similar to human MPM.

**Figure 9.**
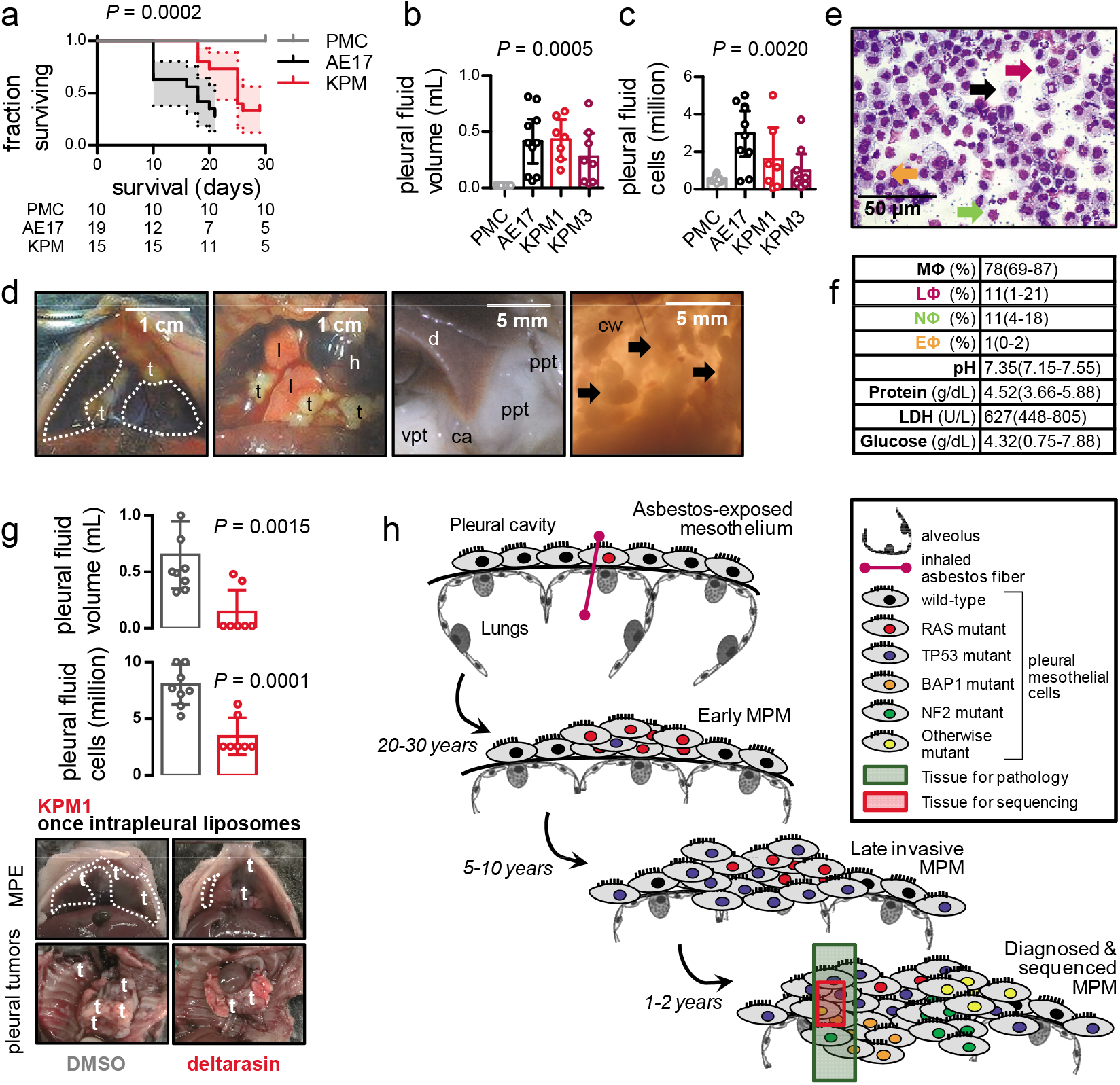
Transplantable and actionable murine mesothelioma models using KPM cells. *C57BL/6* mice received 2 x 10^5^ intrapleural PMC, AE17, or KPM cells. **a,** Kaplan-Meier survival plot. **b,c,** Pleural effusion volume and total cells (*n* = 7-10/group). **d,** Images of the chest before and after opening, showing effusion (dashed lines), visceral (vpt) and parietal (ppt) pleural tumours on the costophrenic angle (ca), the diaphragm (d), and the chest wall (cw). t, tumours; l, lungs; h, heart. **e,** May-Gruenwald-Giemsa-stained pleural cells (macrophages, MΦ: black arrows; lymphocytes, LΦ: purple arrows; neutrophils, NΦ: green arrows; eosinophils, EΦ: orange arrows). **f,** Effusion cytology and biochemistry summary (n = 10). **g,** *C57BL/6* mice received pleural KPM1 cells followed by a single intrapleural injection of liposomes containing 1% DMSO or 15 mg.Kg^−1^ deltarasin in 1% DMSO at day 9 post-tumour cells. Shown are representative images of pleural effusions (dashed lines) and tumours (t), and data summaries of MPE volume (*n* = 7-9/group) and pleural fluid nucleated cells at day 19 post-KPM1 cells. **h,** Proposed role of RAS mutations in mesothelioma. KRAS and other RAS pathway driver mutations initiate some cases of asbestos-induced malignant pleural mesothelioma (MPM), but are lost during tumour evolution, or persist at a clonal level and go undetected during sampling or sequencing. RAS, rat sarcoma; TP53, tumor-related protein 53; BAP1, BRCA1-associated protein 1; NF2, neurofibromatosis 2. Data are presented as survival fraction ± 95% confidence interval (shaded areas) and number of surviving mice (a) or mean ± 95% confidence interval (all other graphs). *P*, overall probability, log-rank test (a), one-way ANOVA (b, c), or unpaired t test (g). ***: *P* < 0.001 for comparison with PMC. LDH, lactate dehydrogenase; ANOVA, analysis of variance.

**Figure 10.**
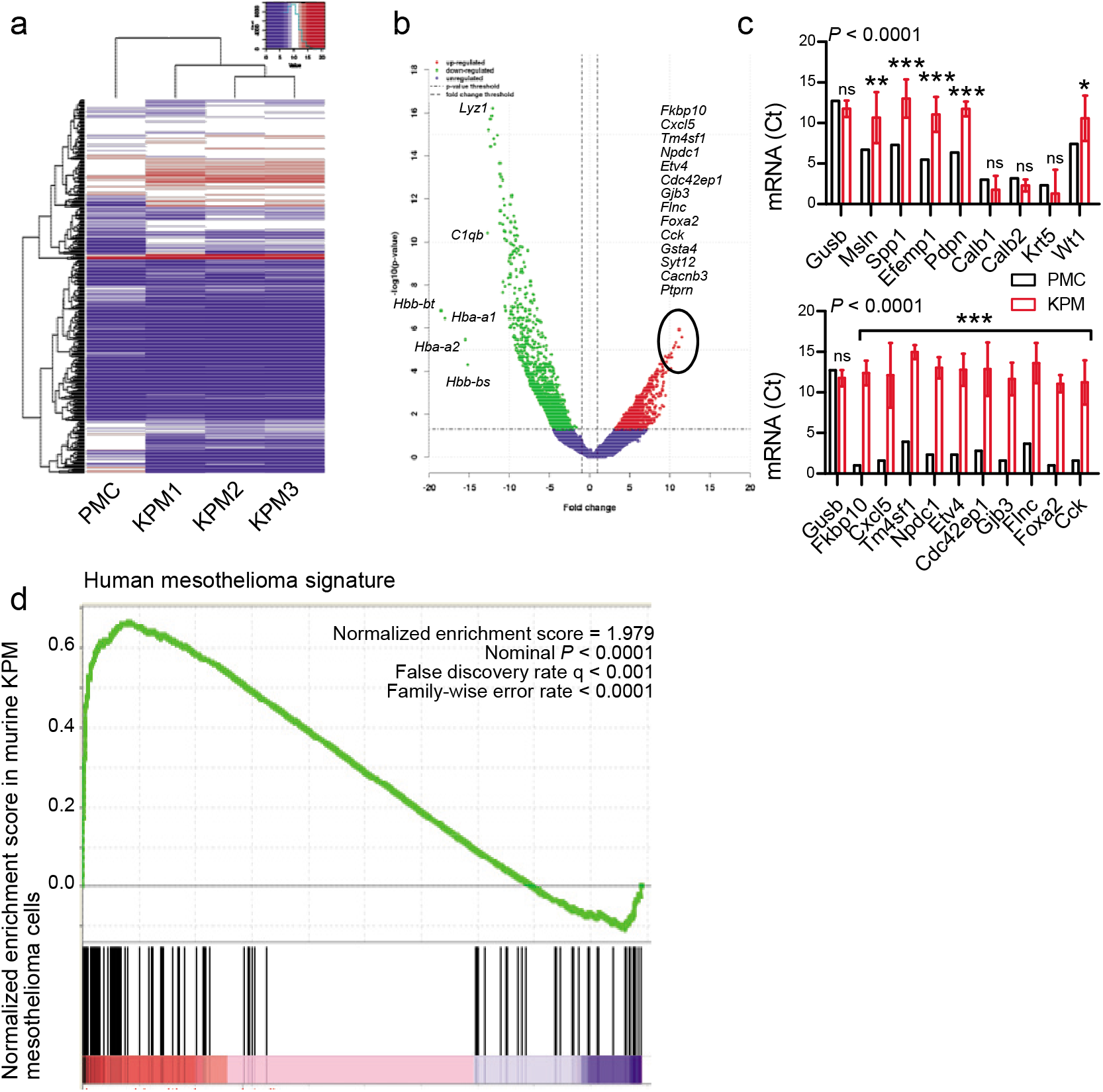
Molecular signature of human mesothelioma in KPM cells. RNA sequencing results (GEO dataset GSE94415) of *KRAS^G12D^;Trp53f/f* mesothelioma (KPM) cells (*n* = 3) compared with pleural mesothelial cells (PMC; *n* = 1 pooled triplicate). a, Unsupervised hierarchical clustering shows distinctive gene expression of KPM versus PMC. b, Volcano plot showing some top KPM versus PMC differentially expressed genes. c, KPM and PMC expression of classic mesothelioma markers (top) and top KPM versus PMC overexpressed genes (bottom). Data given as mean ± 95% confidence interval. *P*: probability, two-way ANOVA. ns, *, **, and ***: *P* > 0.05, *P* < 0.05, *P* < 0.01, and *P* < 0.001 compared with PMC by Bonferroni post-tests. d, Gene set enrichment analysis, including enrichment score and nominal probability value of the 150 gene-signature specifically over-represented in human mesothelioma compared with other thoracic malignancies derived from 113 patients^50^ within the transcriptome of KPM cells versus PMC shows significant enrichment of the human mesothelioma signature in KPM cells.

## DISCUSSION

Our results demonstrate that oncogenic *KRAS* can potentially drive the murine mesothelium towards MPM development. Targeting of oncogenic *KRAS^G12D^* alone to the pleural mesothelium caused epithelioid MPM in mice and together with *Trp53* deletion resulted in biphasic MPM with MPE. We further show that murine MPM carry the initiating *KRAS^G12D^/Trp53* mutations and multiple secondary *Bap1* mutations, are transplantable and druggable, and highly similar to human MPM in terms of molecular markers and gene expression. Finally, we present evidence of frequent *KRAS* point mutations and gains in human MPM. Collectively, the data reveal a potential pathogenic role for *KRAS* mutations in human MPM, and provide multiple new models to study the disease’s pathogenesis and therapy.

Our surprising findings reconcile the paucity of *KRAS* mutations in next generation sequencing studies of MPM^13–16,22^, the sporadic detection of *KRAS* and other related driver mutations (*EGFR, PIK3CA, BRAF*, etc.) in targeted sequencing studies^17–21,23^, and the inefficacy of standalone *Bap1, Cdkn2a, Nf2*, or *Trp53* deletion to cause MPM in mice^31–34^, and support a unifying concept for MPM pathogenesis (Fig. 9h). Mesothelial *KRAS* and other RAS pathway mutations may initiate MPM in some patients, but may be subsequently lost during tumour evolution, or may persist at a clonal level going undetected during sampling/sequencing, as has been shown for lung tumours^51,52^. Indeed, MPM cell lines display RAS pathway activation and *KRAS* mutations^23,35,36^. Importantly, *NF2* is a RAS suppressor^24^, *PIK3CA* mutations were extremely frequent in one study^19^, and the MPM-initiating role ascribed here to *KRAS*^G12D^ is possibly a class effect of RAS pathway mutations, including *NF2* loss and *EGFR*, *BRAF*, *PIK3CA*, and other activating mutations. Taken together, our data and the literature support that *KRAS* and other RAS pathway mutations are present in a subset of tumour cells and patients with MPM.

We also corroborate the critical role of *TP53* in MPM progression. Although standalone *Trp53* deletion did not induce MPM, it promoted *KRAS*^G12D^-driven MPM progression and biphasic histology, as was also observed in combination with *Nf2* but not with *Tsc1* deletion^31,32^, suggesting that *Trp53* loss may selectively cooperate with aberrant RAS signalling in MPM. In addition, *Trp53 loss* rendered *KRAS^G12D^*-driven MPM MPE-competent, a human MPE phenotype reproduced in mice for the first time^31–34^. Again, *Trp53* loss was not causative, but likely potentiated the effusion-promoting effects of mutant *KRAS*, which we recently identified in metastatic effusions^35^. TP53 pathway mutations are frequent in MPM, especially when *CDKN2A*, *BAP1*, and *STK11* mutations are considered as *TP53*-related^25–27^. Taken together with an elegant previous study of combined pleural *Nf2/Trp53* deletion^31^, our findings functionally validate the role of *TP53* mutations in human MPM in driving biphasic histology, tumour progression and metastasis, and poor survival^14,19^. Hence TP53-targeted therapies may be prioritized for biphasic/sarcomatoid MPM subtypes when they become available^53^.

Another surprising finding were the multiple and different *Bap1* mutations of our MPM cell lines, since they originated from tumours inflicted by *KRAS*^G12D^ and *Trp53* loss. Frequent copy number loss and recurrent somatic mutations in *BAP1* have been identified in MPM^13,16,34^. Based on the multiplicity and variety of the *Bap1* mutations we observed, we postulate that they were secondarily triggered by the genomic instability caused from combined *KRAS* mutation and *TP53* loss. Whatever their cause may be, their presence strengthens our findings of a causative role for *KRAS* mutations in MPM, as well as the relevance of the novel mouse models we developed, since *Bap1* is the single most commonly mutated gene in human MPM^13,16,34,54^.

Research on MPM is hampered by the paucity of mouse models of the disease^31^. We provide multiple new mouse models with defined phenotype, histology, and latency: i) a mouse model of pleural epitheliod MPM; ii) genetic and transplantable models of pleural and peritoneal biphasic MPM with accompanying MPE; and iii) three new MPM cell lines of defined genotype, transcriptome, and phenotype syngeneic to *C57BL/6* mice. These contributions are positioned to enhance research in the field by overcoming the need for immune compromise thereby providing intact immune responses critical for MPM^55–58^, by widening the armamentarium of existing cell lines^30,35^, by recapitulating the phenotype of MPM-associated MPE^31–34^, and by experimentally addressing the pleural disease form of mesothelioma^31^.

In conclusion, our findings support that oncogenic *KRAS* causes and *TP53* loss accelerates MPM in mice, and provide new tools for drug discovery. The results are likely applicable to *NF2, PIK3CA, EGFR, BRAF, CDKN2A*, and/or *BAP1*-mutant MPM, since *KRAS*/*TP53* functions likely represent mutation class effects. Many of those are currently druggable, such as *EGFR* and *BRAF* mutations, while novel anti-*KRAS* and *TP53* agents will likely become available soon^53,59–61^. Hence, in addition to revealing a key role for RAS/TP53 pathway mutations in MPM development and progression, we provide unique tools to identify addicted oncogenes and pathways^62^ that will hopefully lead to therapeutic innovations against MPM.

## METHODS

Additional methods are described in the Online Supplement.

### Reagents

Adenoviruses type 5 (Ad) encoding *Melanotus* luciferase (*Luc*) or CRE-recombinase (*Cre*) were from the Vector Development Laboratory, Baylor College of Medicine (Houston, TX), 3-(4,5-dimethylthiazol-2-yl)-2,5-diphenyltetrazolium bromide (MTT) assay from Sigma (St. Louis, MO), and D-luciferin from Gold Biotechnology (St. Louis, MO). Antibodies and primers are listed in Supplementary Tables S2 and S3.

### Humans

The Munich clinical studies were conducted in accordance with the Helsinki Declaration and were prospectively approved by the Ludwig-Maximilians-University Munich Ethics Committee (approvals #623–15 and #711–16). All patients gave written informed consent *a priori*. Pleural fluid was centrifuged at 300 g for 10 min at 4 °C, genomic DNA was extracted using TRIzol (Thermo Fisher Scientific, Waltham, MA) and purified using GenElute Mammalian Genomic DNA Miniprep (Sigma-Aldrich, St. Louis, MO), and 200 ng DNA were used to analyze *KRAS* codon 12/13 and 61 mutations with ddPCR *KRAS* G12/G13 and *KRAS* G61 Screening Kits and QuantaSoft Analysis Pro software (Bio-Rad, Hercules, CA) as described elsewhere^63^. Data were normalized by accepted droplet numbers to yield absolute mutation percentages, which were determined according to the formula:

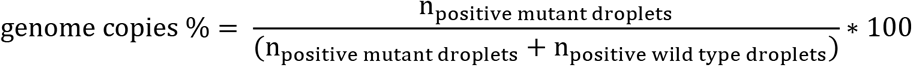

The Nantes clinical studies were conducted in accordance with the Helsinki Declaration and were prospectively approved by the University of Nantes Ethics Committee (approval #xxxx). All patients gave written informed consent *a priori*. MPE samples from 61 patients with MPM were used to generate cell lines, as described elsewhere^39,40^. Genomic DNA from 32 MPM cell lines was extracted with Nucleospin Blood kit (Macherey-Nagel, Düren, Germany) and 500 ng were hybridized to Affymetrix CytoScanHD Arrays (Affymetrix, Santa Clara, CA). Detection, quantification, and visualization of copy number alterations were performed using Affymetrix Chromosome Analysis Suite v3.1.1.27 (Affymetrix, Santa Clara, CA) and data are available at GEO datasets (GSE134349).

### Mice

*C57BL/6* (#000664), *B6.129(Cg)-Gt(ROSA)26Sor^tm4(ACTB-tdTomato,-EGFP)Luo^/J (mT/mG;* #007676)^42^, *FVB-Tg(CAG-luc,-GFP)L2G85Chco/J* (*CAG.Luc.eGFP*; #008450)^64^, B6.129S4-*Kras^tm4Tyj^/J (KRAS^G12D^;* #008179)^44^, and *B6.129P2-Trp53^tm1Brn^/J (Trp53f/f*; #008462)^45^ mice were obtained from Jackson Laboratories (Bar Harbor, ME) and bred on the *C57BL/6* background at the University of Patras Center for Animal Models of Disease. Experiments were approved by the Prefecture of Western Greece’s Veterinary Administration (approval 118018/578-30.04.2014), and were conducted according to Directive 2010/63/EU (http://eur-lex.europa.eu/legal-content/EN/TXT/?uri=CELEX%3A32010L0063). Sex-, weight (20-25 g)-, and age (6-12 week)-matched experimental mice were used, and their numbers are detailed in Supplementary Table S4.

### Mesothelial transgene delivery

Isoflurane-anesthetized *C57BL/6* and *mT/mG* mice received 5 x 10^8^ PFU intrapleural or intraperitoneal Ad-*Cre* or *Ad-Luc* in 100 μL PBS, were serially imaged for bioluminescence on a Xenogen Lumina II after receiving 1 mg retro-orbital D-luciferin under isoflurane anesthesia and data were analyzed using Living Image v.4.2 (Perkin-Elmer, Waltham, MA)^35,65^, or were euthanized and pleural lavage was performed, lungs were explanted, and parietal pleura was stripped. For pleural lavage, 1 mL PBS was injected, was withdrawn after 30 sec, and was cytocentrifuged onto glass slides (5 x 10^4^ cells, 300 g, 10 min) using CellSpin (Tharmac, Marburg, Germany). Lungs were embedded in optimal cutting temperature (OCT; Sakura, Tokyo, Japan) and sectioned into 10 μm cryosections. The parietal pleura was placed apical-side-up onto glass slides. Samples were stained with Hoechst 55238 and were examined on AxioObserver D1 (Zeiss, Jena, Germany) or TCS SP5 (Leica, Heidelberg, Germany) microscopes.

### Primary MPM models

Wild-type (*Wt*), *KRAS^G12D^*, and *Trp53f/f* mice were intercrossed and all possible offspring genotypes received isoflurane anesthesia and 5 x 10^8^ PFU intrapleural or intraperitoneal Ad-*Cre*. Mice were monitored daily and sacrificed when moribund or prematurely for pathology. Mice with pleural fluid volume ≥ 100 μL were judged to have effusions that were aspirated. Animals with pleural fluid volume < 100 μL were judged not to have effusions and underwent pleural lavage. For isolation of primary murine pleural mesothelial cells (PMC), pleural myeloid and lymphoid cells were removed by pleural lavage followed by pleural instilation of 1 mL DMEM, 2% trypsin EDTA, aspiration after 1 min, and culture.

### Transplantable mesothelioma cell lines

Murine *KRAS^G12D^;Trp53f/f* pleural mesotheliomas were minced and cultured in DMEM 10% FBS for > 30 passages, yielding three *KRAS^G12D^;Trp53f/f* mesothelioma (KPM1-3) cell lines, which were compared to AE17 cells (*Kras^G12C^*-mutant asbestos-induced murine mesothelioma)^35^ and PMC. For this, 2 x 10^5^ cells in 100 μL PBS were delivered intrapleurally to isoflurane-anesthetized *C57BL/6* mice that were followed as above. For solid tumour formation, *C57BL/6* mice received 10^6^ subcutaneous PMC, KPM, or AE17 cells in the rear flank, three vertical tumour dimensions (δ^1^, δ^2^, δ^3^) were monitored serially, and tumour volume was calculated as πδ^1^δ^2^δ^3^/6 (29, 34). RNA sequencing was done on an IonTorrent sequencer (Thermo Fisher), data were deposited at GEO datasets (GSE94415), and were analyzed using Bioconductor. Gene set enrichment was done with the Broad Institute pre-ranked GSEA module^66^.

### Liposome preparation and physicochemical characterization

Deltarasin-encapsulating liposomes were prepared as described elsewhere^67,68^, by freeze-drying 30 mg of empty DSPC/PG/Chol (9:1:5 mol/mol/mol) unilamelar sonicated vesicles with 1 mL of deltarasin solution (5 mg.mL^−1^) in PBS, or plain PBS (for empty liposomes), followed by controlled rehydration. Liposome size was decreased by extrusion though Lipo-so-fast extruder polycarbonate membranes (Avestin Europe, Mannheim, Germany) with 400 nm pore diameter. Liposome lipid concentration, size distribution, surface charge (zeta-sizer, Malvern Panalytical Ltd, Malvern, United Kingdom), and drug encapsulation efficiency after measuring the non-liposomal drug absorption at 284 nm, were estimated as reported elsewhere^67,68^.

### *In vivo* drug treatments

Deltarasin-encapsulating liposomes were delivered intrapleurally into *C57BL/6* mice nine days post-intrapleural KPM1 cells, when the first pleural tumors where already established^35^.

### Statistics

Sample size was estimated using G*power^69^ assuming α = 0.05, β = 0.05, and d = 1.5. Animals were allocated to treatments by alternation and transgenic animals casecontrol-wise. Data acquisition was blinded and no data were excluded from analyses. Data were distributed normally by Kolmogorov-Smirnov test and are given as mean ± 95% confidence interval (CI). Sample size (*n*) refers to biological replicates. Differences in means were examined by t-test or one-way analysis of variance (ANOVA) with Bonferroni post-tests and in frequencies by Fischer’s exact or χ^2^ tests. Longitudinal (bioluminescence, MTT, tumour growth) data were analyzed by twoway ANOVA with Bonferroni post-tests. Survival was analyzed using Kaplan-Meier estimates and log-rank test. Probability (*P*) values are two-tailed and *P* < 0.05 was considered significant. Analyses and plots were done on Prism v8.0 (GraphPad, La Jolla, CA).

## END NOTES

### Supplementary information

This paper contains supplementary information pertaining to the methods used and uncropped PCR gels shown.

## Acknowledgements

AE17 cells were a gift from Dr. Y. C. Gary Lee (University of Western Australia, Perth, Australia). This work was supported by European Research Council 2010 Starting Independent Investigator (#260524) and 2015 Proof of Concept Grants (#679345) G. T. S., the Greek State Scholarship Foundation programme “Reinforcement of Postdoctoral Researchers” co-financed by the European Union Social Fund and Greek national funds (NSRF 2014-2020; to I. G.), and a REPSIRE2 European Respiratory Society Fellowship (LTRF 2015-1824; to I. P.).

## Author contributions

A. M. and A. C. K. designed and carried out experiments, analyzed data, provided critical intellectual input, and wrote portions of the paper draft; C. B. and M. G. designed and carried out karyotype analyses, and provided the French MPM cell line cohort; M. A. A. P., A. S. L., C. M. H., I. K., M. L., and R. A. H. designed and carried out sequencing experiments and analysis, droplet PCR, and provided the German MPM tumour cohort; M. I. performed *in vivo* CRE reporter assays; A. C. K. and I. L. performed molecular phenotyping of murine tumours; D. E. W. performed GSEA; H. P. evaluated and diagnosed pathology; S. G. A. prepared liposomes; I. P., M. S., and I. G. designed and performed experiments and provided critical intellectual input and partial funding; and G. T. S. conceived the idea, obtained funding, supervised the study, designed experiments, analyzed data, wrote the paper, and is the guarantor of the study’s integrity. All authors reviewed and concur with the submitted manuscript.

## Correspondence and requests for materials should be addressed to

Georgios T. Stathopoulos, MD PhD; Comprehensive PneumologyCenter, Max-Lebsche-Platz 31, 1. OG, 81377 Munich, Germany, Phone: +49 (89) 3187 1194, Fax: +49 (89) 3187 4661.

## Data availability

Microarray (GEO dataset GSE134349) and RNA sequencing (GEO dataset GSE94415) data generated for this study are freely available at https://www.ncbi.nlm.nih.gov/gds.

## ONLINE SUPPLEMENTARY INFORMATION

## SUPPLEMENTARY METHODS

### Bone marrow transfer

For adoptive BMT, *C57BL/6* mice received 10^7^ bone marrow cells obtained from *CAG.Luc.eGFP* donors i.v. 12 hours after total-body irradiation (1100 Rad). Full bone marrow reconstitution was completed after one month, as described elsewhere^1^.

### PCR and Sanger sequencing

Cellular RNA was isolated using Trizol (Thermo Fisher Scientific, Waltham, MA) followed by RNAeasy purification and genomic DNA removal (Qiagen, Hilden, Germany). For tumor RNA, tissues were passed through 70 μm strainers (BD Biosciences, San Jose, CA) and 10^7^ cells were subjected to RNA extraction. One μg RNA was reverse-transcribed using Oligo(dT)_18_ and Superscript III (Thermo Fisher). Kras, KRAS, Trp53, Bap1, Nf2, Cdkn2a, and Gusb cDNAs were amplified using specific primers (Supplementary Table S2) and Phusion Hot Start Flex polymerase (New England Biolabs, Ipswich, MA). DNA fragments were run on 2% agarose gels or were purified with NucleoSpin gel and PCR clean-up columns (Macherey-Nagel, Düren, Germany) and were sequenced using their primers by VBC Biotech (Vienna, Austria). qPCR was performed using specific primers (Supplementary Table S3) and SYBR FAST qPCR Kit (Kapa Biosystems, Wilmington, MA) in a StepOne cycler (Applied Biosystems, Carlsbad, CA). Ct values from triplicate reactions were analyzed with the 2^−ΔCT^ method^2^. mRNA abundance was determined relative to glycuronidase beta (Gusb) and is given as 2^−ΔCT^ = 2 ^−(Ct of transcript)-(Ct of Gusb)^. The Sanger sequencing trace files were further analyzed for double peak parser using Bioconductor (https://www.bioconductor.org/) with a threshold of 25 Phred quality core^5^. The mismatch basecalls in respect to the wild type were grouped by cell line and used as template to generate the lollipop plot per each KPM cell line for a visual representation of all the mutations detected^6^.

### RNA sequencing

RNA sequencing was done on an IonTorrent sequencer (Thermo Fisher) and data were analyzed using Bioconductor (https://www.bioconductor.org/). File alignments were performed with Tmap(https://github.com/iontorrent/TMAP). Coverage and alignments plot from sequencing were generated using Integrative genome viewer^4^. Alignments are represented as gray polygons with reads mismatching the reference indicated by colour. Loci with a large percentage of mismatches relative to the reference are flagged in the coverage plot as colour-coded bars. Alignments with inferred small insertion or small deletion are represented with vertical or horizontal bars respectively. Gene set enrichment analysis (GSEA) was performed with the Broad Institute pre-ranked GSEA module software (http://software.broadinstitute.org/gsea/index.jsp). The raw bam files, one for each RNA-Seq sample, were summarized to a gene read counts table, using the Bioconductor package GenomicRanges. In the final read counts table, each row represented one gene, each column one RNA-Seq sample and each cell, the corresponding read counts associated with each row and column. The gene counts table was normalized for inherent systematic or experimental biases (e.g. sequencing depth, gene length, GC content bias etc.) using the Bioconductor package DESeq after removing genes that had zero counts over all the RNA-Seq samples (20007 genes). The output of the normalization algorithm was a table with normalized counts, which can be used for differential expression analysis with statistical algorithms developed specifically for count data. Prior to the statistical testing procedure, the gene read counts were filtered for possible artifacts that could affect the subsequent statistical testing procedures. Genes presenting any of the following were excluded from further analysis: i) genes with length less than 500 (2051 genes), ii) genes whose average reads per 100 bp was less than the 25th percentile of the total normalized distribution of average reads per 100bp (0 genes with cutoff value 0.02248 average reads per 100 bp), iii) genes with read counts below the median read counts of the total normalized count distribution (11358 genes with cutoff value 16 normalized read counts). The total number of genes excluded due to the application of gene filters was 5298. The total (unified) number of genes excluded due to the application of all filters was 32595. The resulting gene counts table was subjected to differential expression analysis for the contrast KPM versus PMC using the Bioconductor package DESeq. The final numbers of differentially expressed genes were 2344 statistically significant genes were found and of these, 650 were up-regulated and 1694 were down-regulated according to an absolute fold change cutoff value of 2.

### Cell and tissue analyses

MPE fluid was diluted in ten-fold excess red blood cells lysis buffer (155 mM NH_4_Cl, 12 mM NaHCO_3_, 0.1 mM EDTA). Total pleural cell counts were determined microscopically in a haemocytometer and cytocentrifugal specimens (5 x 10^4^ cells each) of pleural fluid cells were fixed with methanol for 2 min. Cells were stained with May-Grünwald stain in 1 mM Na_2_HPO_4_, 2.5 mM KH_2_PO_4_, pH = 6.4 for 6 min and Giemsa stain in 2 mM Na_2_HPO_4_, 5 mM KH_2_PO_4_, pH = 6.4 for 40 min, washed with H_2_O, and dried. Slides were mounted with Entellan (Merck Millipore, Darmstadt, Germany), coverslipped, and analyzed. For flow cytometry, 10^6^ nucleated pleural fluid cells suspended in 50 μL PBS supplemented with 2% FBS and 0.1% NaN_3_ were stained with the indicated antibodies according to manufacturer’s instructions (Supplementary Table S2) for 20 min in the dark, washed, and resuspended in buffer for further analysis. Lungs, visceral pleural tumors, parietal pleural tumors, and chest walls were fixed in 4% paraformaldehyde overnight, embedded in paraffin or OCT and were stored at room temperature or −80°C, respectively. Five-μm paraffin or 10-μm-cryosections were mounted on glass slides. Sections were labeled using the indicated antibodies (Supplementary Table S2), counterstained with Envision (Dako, Carpinteria, CA) or Hoechst 33258 (Sigma), and mounted with Entellan new (Merck Millipore) or Mowiol 4-88 (Calbiochem, Gibbstown, NJ). For isotype control, primary antibody was omitted. Bright-field and fluorescent microscopy was done on AxioLab.A1 (Zeiss), AxioObserver.D1 (Zeiss), or TCS SP5 (Leica) microscopes and digital images were processed with Fiji^3^.

### Cell culture

All KPM cell lines were deposited at the Laboratory for Molecular Respiratory Carcinogenesis cell line facility and are available upon request. Cells were cultured at 37°C in 5% CO_2_-95% air using DMEM 10% FBS, 2 mM L-glutamine, 1 mM pyruvate, 100 U/ml penicillin, and 100 mg/ml streptomycin and were tested biannually for identity (by short tandem repeats) and *Mycoplasma Spp*. (by PCR). For *in vivo* injections, cells were harvested with trypsin, incubated with Trypan blue, counted on a hemocytometer, and > 95% viable cells were injected into the pleural space (2 x 10^5^) or into the skin (10^6^) as described elsewhere^1^. *In vitro* cell proliferation was determined using MTT assay.

## SUPPLEMENTARY FIGURES

**Supplementary Figure S1.**
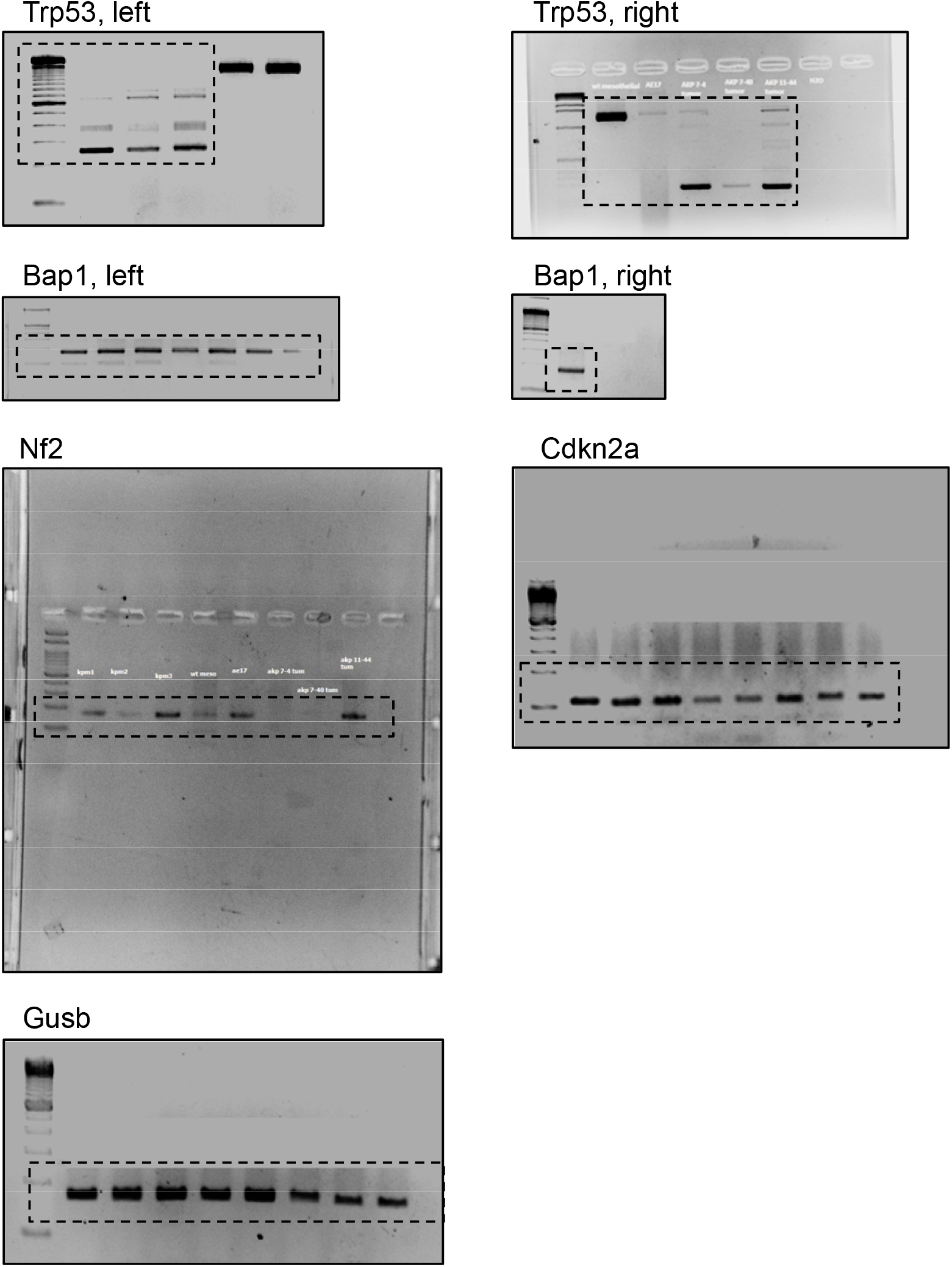
Uncropped PCR gels from Figure 7f. Cropboxes indicate the gel areas shown in the main Figure.

## SUPPLEMENTARY TABLES

**Supplementary Table S1.**
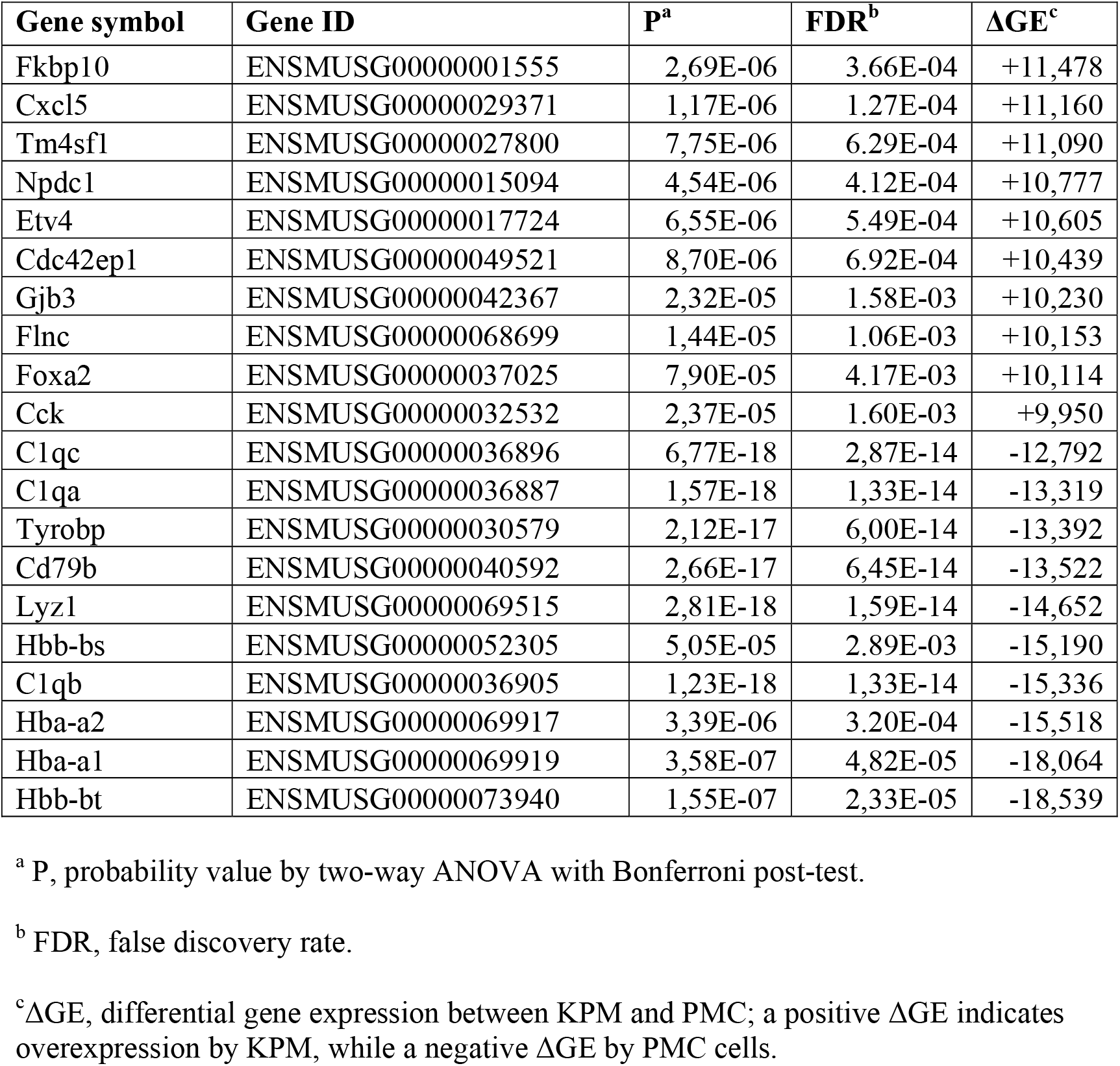
Top differentially expressed genes in *KRASG12D;Trp53f/f* malignant pleural mesothelioma cell lines (KPM) versus pleural mesothelial cells (PMC).

**Supplementary Table S2.** Antibodies used for these studies.

| **Method<sup>a</sup>** | **Target<sup>b</sup>** | **Provider<sup>c</sup>** | **Catalog #** | **Dilution** | **Conjugate<sup>d</sup>** |
| --- | --- | --- | --- | --- | --- |
| IHC | PCNA | Abcam | ab2426 | 1:2000 | - |
|  | CAL | Abcam | ab203055 | 1:400 | - |
|  | PDPN | Merck | ABT34 | 1:1000 | - |
|  | OPN | Abcam | ab8448 | 1:100 | - |
|  | KRT5/6 | Merck | MAB3412 | 1:200 | - |
|  | SFTPC | Santa Cruz | FL-197 | 1:100 | - |
|  | BAP1 | Santa Cruz | Sc-28383 | 1:100 | - |
| FC | CD11b | eBioscience | 12-0112 | 0.1µg/10<sup>6</sup>cells | PE |
|  | Gr1 | eBioscience | 25-5931-82 | 0.1µg/10<sup>6</sup>cells | PE-Cy7 |

<sup>a</sup>Application: IHC, immunohistochemistry; FC, flow cytometry.
<sup>b</sup>Target: PCNA, proliferating cell nuclear antigen; CAL, calretinin; PDPN, podoplanin; OPN, osteopontin; KRT, cytokeratin; SFTPC, surfactant protein C.
<sup>c</sup>Providers: Abcam, Cambridge, UK; eBioscience, San Diego, CA; Santa Cruz Biotechnology, Santa Cruz, CA; Merck Millipore, Darmstadt, Germany.
<sup>d</sup>Conjugates: PE, phycoerythrin; PE-Cy7, phycoerythrin-cyanin 7.

**Supplementary Table S3.**
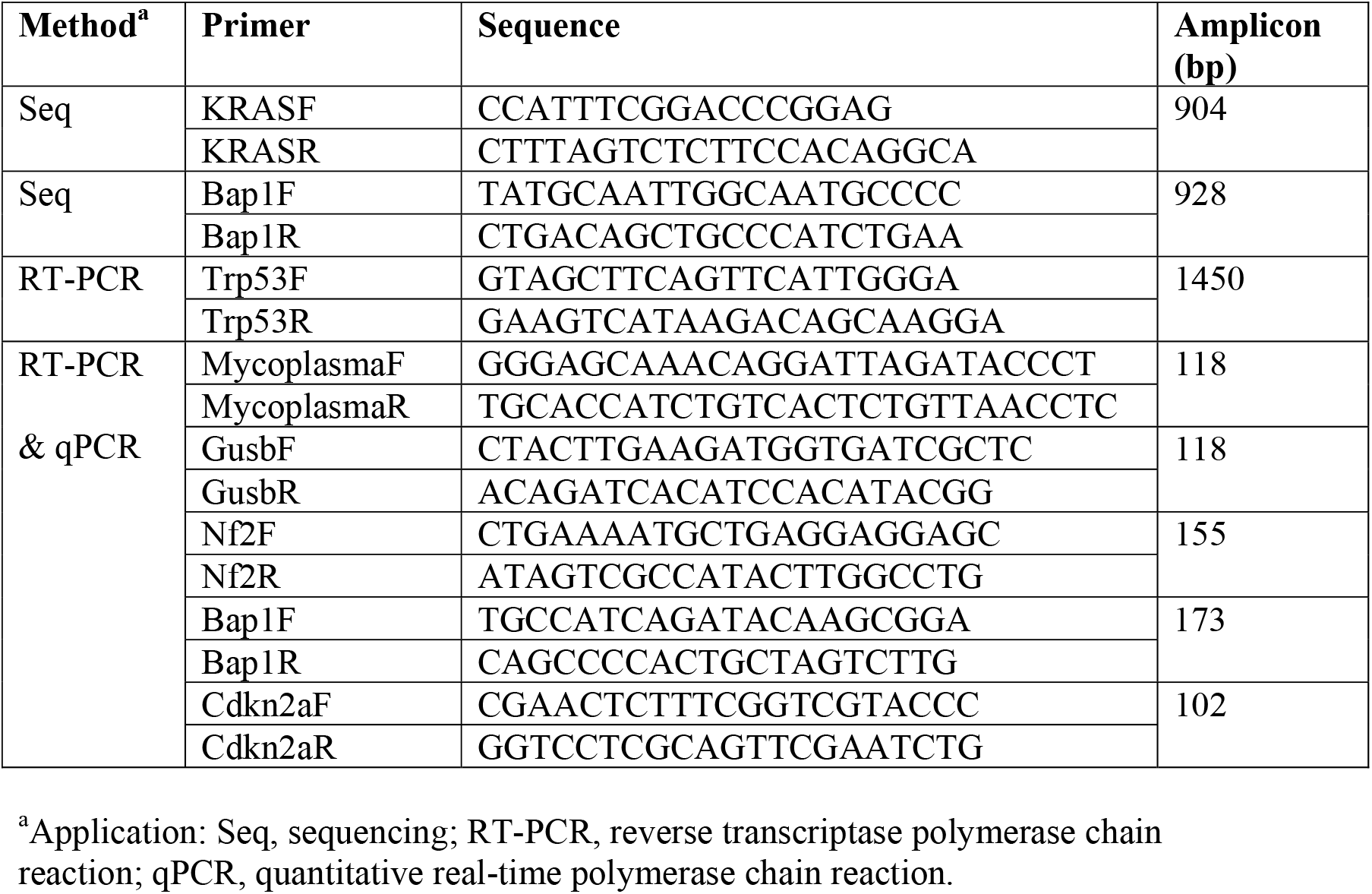
PCR primers used for these studies.

**Supplementary Table S4.** Number of experimental mice (*n*) used for these studies.

| Strain designation | Jackson Laboratory ID # | Short strain designation | *n* |
| --- | --- | --- | --- |
| *C57BL/6J* | 000664 | *C57BL/6* | 177 |
| B6.129(Cg)-Gt(ROSA)26Sor<sup>*tm4*(*ACTB*-*tdTomato*,-EGFP)*Luo*/J</sup> | 007676 | *mT/mG* | 40 |
| B6.129S4-*Kras*<sup>*tm4Tyj*</sup>/J | 008179 | *KRAS*<sup>*G12D*</sup> | 40 |
| B6.129P2-*Trp53*<sup>*tm1Brn*</sup>/J | 008462 | *Trp53f/f* | 33 |
| FVB-Tg(CAG-luc,-GFP)L2G85Chco/J | 008450 | CAG.Luc.eGFP | 5 |
| Intercrossed mice |  | *Trp53f/Wt* | 11 |
|  |  | *KRAS*<sup>*G12D*</sup>; *Trp53f/Wt* | 11 |
|  |  | *KRAS*<sup>*G12D*</sup>; *Trp53f/f* | 115 |
| **Total** |  |  | **432** |

